# Multi-ancestry GWAS reveals loci linked to human variation in LINE-1- and Alu-insertion numbers

**DOI:** 10.1101/2024.09.10.612283

**Authors:** Juan I. Bravo, Lucia Zhang, Bérénice A. Benayoun

**Author notes:** Corresponding author. Leonard Davis School of Gerontology, University of Southern California, Los Angeles, CA, 90089, USA. *E-mail address* (B.A. Benayoun).

## Abstract

LINE-1 (L1) and Alu are two families of transposable elements (TEs) occupying ∼17% and ∼11% of the human genome, respectively. Though only a small fraction of L1 copies is able to produce the machinery to mobilize autonomously, Alu and degenerate L1s can hijack their functional machinery and mobilize *in trans*. The expression and subsequent mobilization of L1 and Alu can exert pathological effects on their hosts. These features have made them promising focus subjects in studies of aging where they can become active. However, mechanisms regulating TE activity are incompletely characterized, especially in diverse human populations. To address these gaps, we leveraged genomic data from the 1000 Genomes Project to carry out a trans-ethnic GWAS of L1/Alu insertion singletons. These are rare, recently acquired insertions observed in only one person and which we used as proxies for variation in L1/Alu insertion numbers. Our approach identified SNVs in genomic regions containing genes with potential and known TE regulatory properties, and it enriched for SNVs in regions containing known regulators of L1 expression. Moreover, we identified reference TE copies and structural variants that associated with L1/Alu singletons, suggesting their potential contribution to TE insertion number variation. Finally, a transcriptional analysis of lymphoblastoid cells highlighted potential cell cycle alterations in a subset of samples harboring L1/Alu singletons. Collectively, our results suggest that known TE regulatory mechanisms may be active in diverse human populations, expand the list of loci implicated in TE insertion number variability, and reinforce links between TEs and disease.

## 1. INTRODUCTION

In the human genome, the two most abundant families of transposable elements (TEs) are Long INterspersed Element-1 (LINE-1; L1) and Alu, which account for ∼16– 17% and ∼9–11% of the genome, respectively [1, 2]. Full-length L1 elements span ∼6 kilobases and produce bicistronic messenger ribonucleic acids (mRNAs) encoding two polypeptides, ORF1p and ORF2p, necessary for L1 transposition (reviewed in [3]). The L1 family can be segregated into 3 subfamilies depending on the evolutionary age of the copy: the L1M (mammalian-wide) lineage is the oldest, the L1P (primate-specific) lineage is of intermediate age, and the L1PA lineage is the youngest. Importantly, only the L1PA1/L1Hs subfamily contains ∼80-100 actively mobile copies in the average human genome [4], with the remaining ∼500,000 L1 copies being rendered non-autonomous due to the presence of loss-of-function mutations or truncations [1]. In contrast to L1 elements, Alu elements are short (∼300 bp) non-autonomous retrotransposons that rely on functional L1 machinery for their mobilization [5–7]. Alu retrotransposons can also be segregated by evolutionary age into the following subfamilies: AluJ is the oldest lineage and is likely completely inactive in humans, AluS is the middle-aged lineage and contains mobile copies, and AluY is the youngest lineage and contains the largest number of functionally intact elements [8].

For an expansion of L1 insertions to occur, L1 must undergo a multi-step and tightly regulated lifecycle. This lifecycle begins with transcription of an active, full-length L1 copy and ends with reverse transcription and integration of L1 by a process called target primed reverse transcription (TPRT) (reviewed in [3]). Importantly, though neither Alu elements or degenerate L1 copies can mobilize autonomously, they can hijack proteins from transposition-competent L1s and mobilize *in trans* [6, 7, 9]. Though not traditionally considered part of the L1/Alu lifecycles, other genome remodeling (i.e. recombination or DNA repair) mechanisms can further contribute to TE insertion number variation. This includes, for example, repeat-mediated deletion (RMD) events whereby two repetitive elements (often Alu elements) on the same chromatid recombine and cause the deletion of one of the repeats as well as the intervening sequence, which may include additional repeats [10, 11]. More broadly, non-allelic homologous recombination (NAHR) events ([12–15] and reviewed in [16, 17]) can directly generate large chromosomal deletions and duplications, which may include repetitive sequences. Ultimately, TE insertion number variation is shaped by a combination of *de novo* insertions resulting from their lifecycle and genome remodeling that can expand or retract the number of insertions.

Characterizing the mechanisms governing L1 and Alu transcriptional and mobilization control will be important, given their associations with, and potential contributions to, aging and aging-associated diseases like cancer (discussed in [18–20]). Fundamentally, L1 and/or Alu can alter several hallmarks of aging [21], such as genomic instability, cellular senescence, and inflammation. Though the origin of the signal is unclear (genomic, extra-chromosomal, or cytosolic), an increase in L1 copies has been observed with chronological aging [22] and during cellular senescence [23]. Moreover, a key feature of cellular senescence is the senescence-associated secretory phenotype (SASP) whereby cells secrete an amalgamation of pro-inflammatory factors [24] that may contribute to chronic, low-grade, sterile inflammation with chronological age (a phenomenon referred to as “inflamm-aging”) [24, 25]. L1 can induce a senescent-like state *in vitro* in several cell lines [26, 27] and its cytoplasmic complementary DNA (cDNA) is implicated in the maturation of the SASP response and the establishment of deep senescence through the production of interferons [28]. Similarly, Alu RNA can upregulate senescence markers in retinal pigment epithelium (RPE) cells from human eyes with geographic atrophy [29], and knockdown of Alu transcripts was reported to promote senescence exit in adult adipose-derived mesenchymal stem cells [30]. These findings highlight the relevance of L1 and Alu retrotransposons in pathological, age-associated features and highlight the important of characterizing TE control mechanisms.

To maintain homeostasis, it is imperative that host cells tightly regulate TE activity (reviewed in [31, 32]). These mechanisms, however, remain incompletely characterized due to the cell-specific and transposon-specific nature of TE regulatory mechanisms. Indeed, no systematic, genome-wide screen for regulators of Alu expression or mobilization has been carried out thus far, to our knowledge, and our understanding of L1 control mechanisms remains incomplete, limiting our ability to understand why they are derepressed during aging. To address these gaps, a number of *in vitro* and *in silico* approaches have been developed to scan for novel regulators of TE expression or mobilization. *In vitro* approaches have relied on clustered regularly interspaced short palindromic repeats (CRISPR)-based and small RNA-based tools to decipher L1 regulation in several types of cancer cells [33–37]. These approaches, however, can be technically challenging to implement in non-cancerous cells, like primary cells, which may not tolerate hyper-elevated transposon activity or that may resist genetic perturbations. To complement these methods, a number of *in silico* approaches have been developed that utilize chromatin immunoprecipitation followed by sequencing (ChIP-seq) [38], gene co-expression networks [39], or insertion number-expression correlations [40] to explore L1 regulation without external manipulations. More recently, we screened for candidate regulators of L1 RNA levels in lymphoblastoid cell lines (LCLs) using trans-expression quantitative trait locus (trans-eQTL) analysis [41]. These tools highlight the need for, and usefulness of, alternative approaches that utilize increasingly available, large ‘-omic’ datasets to identify potentially novel mechanisms of TE control.

In this study, we develop a genome-wide association study (GWAS) pipeline to identify genomic loci associated with variation in global L1 and Alu insertion singletons in diverse human populations. Global singleton insertions reflect rare, recently acquired insertions observed uniquely in only one person in a population [42]. Thus, we used insertion singletons as proxies for L1/Alu insertion number variation, which can arise through *de novo* transposition events or genome remodeling mechanisms. We demonstrate that our GWAS approach captures, and enriches, genomic regions containing known and potential regulators of TE activity. We observe that this approach also captures reference insertions and polymorphic structural variants that may influence L1 or Alu insertion number variation. Finally, we note that associated loci fall into a few genes with clinical relevance, strengthening the association between TEs and disease.

## 2. RESULTS

### 2.1 Identification of genomic loci associated with L1/Alu singletons in diverse human populations

To unbiasedly identify potential genetic sources of L1 and Alu insertion number variation in human populations, we leveraged a publicly available human “omic” dataset with thoroughly characterized genetic information. For this analysis, we utilized 2503 multi-ethnic samples from the 1000 Genomes Project for which both single nucleotide variant (SNV) and structural variant (SV) data were available. Specifically, this included individuals from 5 super-populations: 660 African (AFR), 504 East Asian (EAS), 503 European (EUR), 489 South Asian (SAS), and 347 Admixed American (AMR) individuals who declared themselves healthy at the time of sample collection (**Figure 1A**). As a quality control step, we checked whether the combined SNV and SV data segregated samples by population following principal component analysis (PCA). These analyses demonstrated that the top four principal components segregated population groups within each super-population (**Figure S1A-S1E**).

**Fig 1.**
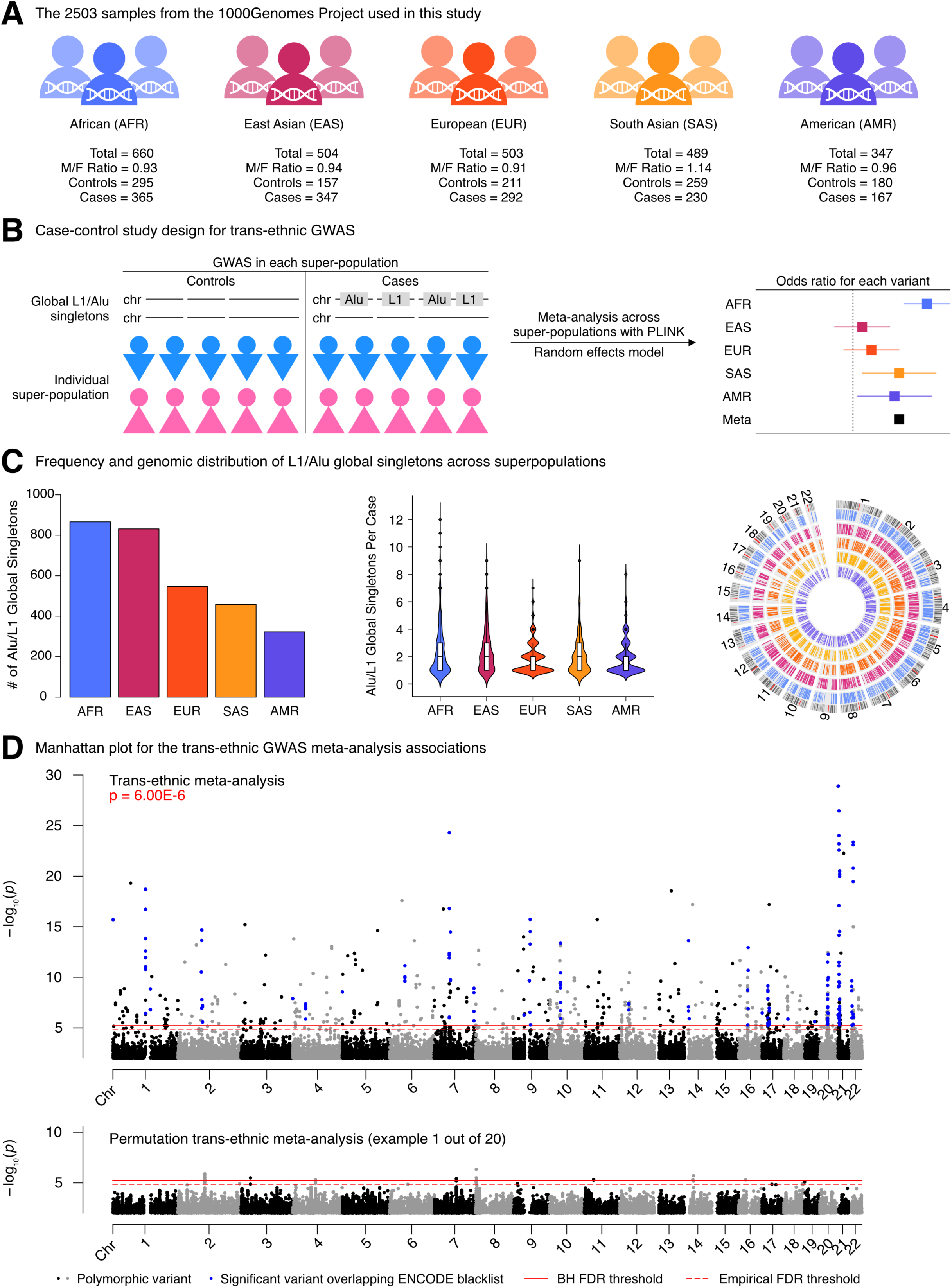
Overview of the pipeline to scan for genetic variants associated with L1/Alu global singletons. **(A)** An illustration of the samples used in this study. SNV and SV genetic data was available for 2503 individuals from 5 super-populations, including 660 Africans (AFR), 504 East Asians (EAS), 503 Europeans (EUR), 489 South Asians (SAS), and 347 Admixed Americans (AMR). Males and females were approximately equally represented, with male-to-female ratios (M/F ratios) ranging from 0.91 to 1.14. **(B)** A schematic illustrating the trans-ethnic integration of available SNV and SV data to identify variants associated with L1/Alu insertion global singletons. Within each super-population, samples were segregated into cases and controls depending on whether or not they harbored a global Alu or L1 insertion singleton. GWAS was carried out within each super-population to identify polymorphic SNVs and SVs associated with case-control status. Finally, GWAS results from all 5 super-populations were meta-analyzed using a random effects statistical model, yielding a summary meta-analysis odds ratio and p-value for each variant. **(C)** The frequency of Alu and L1 insertion singletons in each super-population (*left panel*) or among cases within each super-population (*middle panel*). The distribution of insertion singletons across autosomes is also shown (*right panel*). **(D)** A Manhattan plot for the trans-ethnic GWAS meta-analysis. The dashed line at p = 1.40E-5 corresponds to an average empirical FDR < 0.05, based on 20 random permutations. One such permutation is shown in the bottom panel for illustrative purposes. The solid line at p = 6.00E-6 corresponds to a Benjamini-Hochberg FDR < 0.05. The stricter of the two thresholds, p = 6.00E-6, was used to define significant SNVs and SVs. Significant variants overlapping regions in the ENCODE blacklist v2 are shown in blue and were omitted from downstream analyses. FDR: False Discovery Rate.

We then carried out an integration of the available multi-ethnic SNV and SV genomic data (**Figure 1B**). For the phenotype, we focused on global singleton SVs, which are rare SVs that are observed exactly once in a single person [42], for L1 and Alu insertions. The rarity of these insertions is similar to, but more stringent than, the threshold used in a study of *Arabidopsis* that explored the genetic basis of variable transposition rate [43]. We employed this more stringent threshold (observed in only 1 individual) to increase the likelihood of capturing a recent mobilization event (likely to have occurred in the germline or the early embryonic stage). Unique, or private, insertions are likely to be family-specific rather than population-specific and suggest a recent relaxation of transposon control. For all these reasons, we hypothesized that L1/Alu insertion singletons could serve as proxies for elevated L1/Alu insertion number variation.

Thus, we first split our samples into cases and controls, depending on whether or not they contained an L1 and/or an Alu insertion singleton (**Supplementary Table S1A**). Second, we carried out a GWAS within each super-population to identify common, polymorphic SNVs and SVs associated with case-control status. Third, to maximize statistical power and identify shared, trans-ethnic sources of TE singleton number variation, we meta-analyzed our GWAS results across the 5 super-populations using a random effects statistical model, which allows for increased generalizability across diverse human cohorts compared to fixed effects models [44, 45]. Interestingly, several hundred L1/Alu global singleton insertions were detected in each super-population, ranging from 322 in the American cohort to 866 in the African cohort (**Figure 1C**). Though most case samples had 1-3 L1/Alu global singletons, several samples exhibited much more extreme TE singleton accumulation, especially within the African super-population (**Figure 1C**). Finally, we note that L1/Alu global singleton insertions were distributed across all autosomes in all 5 super-populations (**Figure 1C**). Many L1/Alu singletons resided in intronic (907/3024 ∼ 30%) and intergenic (834/3024 ∼ 27.6%) regions and were enriched for these features compared to background (**Table 1**). This is partially consistent with a prior report where *de novo* engineered L1 insertions were not enriched in genes and endogenous L1 elements were depleted from genic regions in one cell line [46], though the preferences for introns and exons in that study are unclear. Theoretically, the deleterious effects of insertions within intronic and intergenic regions may be blunted compared to the effects of insertions within exonic regions, and we speculate that this may be part of the reason for L1/Alu insertion site preferences. In contrast, L1/Alu singletons were depleted in regions already containing LINE or SINE transposons, compared to their background distribution (**Table 1**). We speculate that the target motif needed for L1/Alu mobilization may be disrupted in these regions, restricting L1/Alu integration into these sites.

**Table 1.** Enrichment of genomic features overlapping global L1/Alu singletons.

| HOMER Annotation | Number of peaks | Log<sub>2</sub> Ratio (obs/exp) | LogP enrichment (+values depleted) |
| --- | --- | --- | --- |
| Intron | 907 | 0.451 | -56.155 |
| Intergenic | 834 | 0.157 | -8.634 |
| DNA | 133 | 0.426 | -7.538 |
| Unknown | 4 | 2.474 | -5.062 |
| snRNA | 2 | 2.681 | -3.229 |
| 3UTR | 35 | 0.371 | -2.529 |
| Low_complexity | 8 | 0.563 | -1.715 |
| TTS | 37 | 0.184 | -1.429 |
| LTR | 261 | 0.042 | -1.142 |
| snoRNA | 0 | -0.001 | 0 |
| SINE? | 0 | -0.004 | 0.003 |
| RC? | 0 | -0.082 | 0.059 |
| tRNA | 0 | -0.127 | 0.092 |
| miRNA | 0 | -0.139 | 0.101 |
| RNA | 0 | -0.149 | 0.109 |
| scRNA | 0 | -0.166 | 0.122 |
| rRNA | 0 | -0.265 | 0.201 |
| srpRNA | 0 | -0.339 | 0.265 |
| RC | 0 | -0.449 | 0.365 |
| DNA? | 0 | -0.506 | 0.42 |
| pseudo | 2 | -0.088 | 0.442 |
| ncRNA | 7 | -0.033 | 0.554 |
| Retroposon | 3 | -0.427 | 0.852 |
| LTR? | 0 | -1.098 | 1.141 |
| 5UTR | 1 | -1.382 | 1.324 |
| Promoter | 30 | -0.267 | 1.746 |
| CpG-Island | 3 | -1.585 | 3.86 |
| Simple_repeat | 22 | -0.615 | 3.863 |
| LINE | 550 | -0.139 | 5.176 |
| Exon | 18 | -1.057 | 8.089 |
| Satellite | 26 | -1.493 | 22.78 |
| SINE | 141 | -1.431 | 112.799 |

As expected, GWAS in each super-population was generally underpowered (**Figure S2A-S2E**). Though we were able to identify many significant (FDR < 0.05) variants in the African cohort (**Figure S2A**), we could not identify significant variants in the East Asian (**Figure S2B**) and European (**Figure S2C**) cohorts, and we were only able to identify a handful of significant variants in the South Asian (**Figure S2D**) and Admixed American (**Figure S2E**) cohorts. These observations, and the abundant associations observed in the African super-population in particular, may be related to the high genetic diversity among African populations [47, 48], including containing more mobile element variants and more population-specific mobile element variants compared to other super-populations [49–51]. In contrast to the individual analyses, the GWAS meta-analysis integrating all super-populations identified 658 significant variants distributed across all 22 autosomes, though there was especially strong and recurrent signal on chromosome 21 (**Figure 1D, Table S1B**). Importantly, to ensure that associations were not completely dependent on the African super-population, we also ran the meta-analysis using only the four non-African super-populations (**Figure S3A-S3C**). Though this analysis had a smaller sample size, we identified 194 significant variants that were shared with the complete meta-analysis and 21 variants that were unique (**Figure S3C, Table S1C**), suggesting that the variants identified in the complete meta-analysis are robust. Moreover, there was significant (p = 5.26E-213) overlap between the variants identified in the non-African meta-analysis and the African-only analysis (**Figure S3D**), further demonstrating that i) the genomic architecture underlying L1/Alu singleton variation is similar between non-African and African super-populations and ii) a subset of variants can be independently identified in two separate populations. Since super-population-specific variants were limited, we focused on the results from the complete meta-analysis. To simplify functional annotation of significant variants and discard potential false positives, we omitted from downstream analyses significant variants overlapping the “ENCODE blacklist v2” [52]. After filtering 188 blacklisted SNVs, we were left with 449 greenlisted SNVs and 21 greenlisted SVs that were significantly associated with case-control status (**Figure 1D**, **Table 2, Supplementary Table S1B**).

**Table 2.** The top 5 most significant, greenlisted variants with an odds ratio > 1 (top) or < 1 (bottom).

| Variant | Chr | Position | Nearby_Genes | Odds_Ratio | P_Meta | P_AFR | P_AMR | P_EAS | P_EUR | P_SAS |
| --- | --- | --- | --- | --- | --- | --- | --- | --- | --- | --- |
| INV_delly_INV00066128 | 21 | 26001780 | ATP5J, APP | 4.3813 | 5.67E-23 | 2.16E-14 | 9.90E-13 | 3.27E-06 | 3.14E-06 | 3.38E-10 |
| ALU_umary_ALU_7919 | 10 | 42173057 | ZNF33B | 3.1524 | 7.75E-14 | 2.42E-07 | 0.000214 | 0.000959 | 0.000269 | 1.61E-08 |
| L1_umary_LINE1_1066 | 5 | 20834958 | CDH18 | 2.2789 | 7.64E-13 | 2.41E-06 | 0.03278 | 0.01845 | 0.03176 | 1.86E-05 |
| SVA_umary_SVA_602 | 14 | 64980975 | RAB15, FNTB | 2.221 | 3.07E-10 | 1.08E-06 | 0.008247 | 0.0006175 | 0.05464 | 0.001859 |
| ALU_umary_ALU_3176 | 4 | 29731532 | PCDH7 | 3.3541 | 5.17E-10 | 2.76E-10 | 0.0006186 | 0.0001465 | 0.0003296 | 4.82E-16 |
| rs199980215 | 1 | 68007166 | GNG12, DIRAS3 | 0.3235 | 4.91E-20 | 1.50E-08 | 5.24E-07 | 0.002965 | 0.03624 | 3.54E-05 |
| rs1317852644 | 13 | 64517674 |  | 0.2899 | 2.87E-19 | 0.02896 | 5.08E-07 | 0.0002641 | 0.00253 | 3.15E-08 |
| rs201291405 | 6 | 47190021 | ADGRF1, TNFRSF21 | 0.2706 | 2.62E-18 | 3.54E-06 | 3.31E-05 | 0.02904 | 0.0103 | 1.36E-07 |
| rs1056816611 | 14 | 31879290 | ARHGAP5, NUBPL | 0.3132 | 6.39E-18 | 1.86E-06 | 0.004238 | 0.01085 | 0.0001221 | 4.84E-07 |
| rs1255393055 | 14 | 31879311 | ARHGAP5, NUBPL | 0.3132 | 6.39E-18 | 1.86E-06 | 0.004238 | 0.01085 | 0.0001221 | 4.84E-07 |

To assess the potential functions of greenlisted, significant variants, nearby genes were assigned to variants and over-representation analysis (ORA) was carried out (**Figure S4A**). Except for 51 greenlisted SNVs that were not linked to any gene, the remaining SNVs were linked to 1-2 genes and were found within 1000 kilobases of a transcriptional start site (TSS) (**Figure S4B**). This observation highlights the association of intergenic and gene-proximal, rather than distal, genetic variation with L1/Alu insertion number differences. Over-representation analysis of the associated genes using the Gene Ontology (GO) Biological Process gene set revealed an enrichment of terms related to heart development (such as ‘regulation of heart growth’, ‘cardiac chamber morphogenesis’, and ‘positive regulation of cardiac muscle cell proliferation’) and neuronal function (such as ‘neuron recognition’ and ‘axonal fasciculation’; **Figure S4B**, **Table S1D**). Interestingly, genes related to ‘reproduction’ were also significantly over-represented among our list of associated genes (**Supplementary Table S1D**). Similar to the SNVs, greenlisted SVs were all linked to 1-2 genes and were within 1000 kilobases of an annotated TSS (**Figure S4C**). Likely due to the low number of greenlisted SVs, and consequently low number of associated genes, we were unable to identify any significantly enriched GO Biological Process gene sets (**Supplementary Table S1E**). Given the limited number of greenlisted SVs and the unavailability of SV sequences, we largely focused on greenlisted SNVs in downstream enrichment analyses.

As a complementary approach, we also predicted the functional impact of significant variants using SnpEff [53] (**Supplementary Table S1F**). Most variants were assigned a ‘modifier’ impact by SnpEff—this annotation describes non-coding variants where definitive functional predictions are not straightforward. Nonetheless, we highlight a few variants which overlapped clinically relevant genes. For example, the most significant, greenlisted variant we identified was an inversion SV (INV_delly_INV00066128, p = 5.67E-23, odds ratio = 4.38) residing in an intronic Alu copy within the *APP* (amyloid beta precursor protein) gene, an important biological marker for Alzheimer’s disease (AD) [54]. Similarly, we identified several SNVs (rs61994687, p = 4.11E-7, odds ratio = 0.54; rs1175403595, p = 3.85E-8, odds ratio = 0.44; rs1343402870, p = 4.49E-6, odds ratio = 0.43) in intronic or downstream regions of *PWRN1* within the Prader-Willi syndrome (PWS) region. To explore non-protein-coding roles greenlisted SNVs may play, we assigned them to ENCODE candidate cis-Regulatory Elements (cCREs) [55] (**Figure S4D**). Although about 8% (36/449) of greenlisted SNVs resided in an ENCODE cCRE, these were significantly depleted (FDR = 6.86E-15) in our greenlisted SNVs compared to background SNVs (**Figure S4D**). Ultimately, our results suggest that proximal, intergenic variation is associated with L1/Alu insertion singleton number variation.

To further explore the potential functional roles of SNV-associated genes, we leveraged expression data from the Genotype-Tissue Expression (GTEx) Portal [56–58] to assess the patterns of tissue expression of SNV-linked genes (**Figure S4E**, **Supplementary Table S1G**). In particular, expression patterns in the brain and gonads were of special interest, given that L1 activity tends to be more frequent in those tissues compared to others (discussed in [59]), and that L1/Alu integration events observed in our GWAS would have to occur in the germline for transmission across generations. Thus, we reasoned that if SNV-associated genes played roles in L1/Alu singleton number variation, they may be more abundantly expressed in the brain and in gonads. Interestingly, there was a cluster of genes that was very abundantly expressed in testes but not in other tissue types (including ovarian tissue), suggesting the existence of potential sex-specific mechanisms of *de novo* L1/Alu insertion transmission. Furthermore, there was also a cluster of genes that were abundantly expressed across brain regions and much less abundantly expressed across other tissue types. More generally, there were many SNV-linked genes that were abundantly expressed in more than ∼50% of tissue types. These results highlight that significant SNV-associated genes have tissue-specific expression patterns, including some genes that are very abundant in tissues with documented L1 activity.

### 2.2 Significant SNVs are enriched near regulators of L1 expression

One of the primary motivations for carrying out this study was to identify novel, candidate regulators of L1 and/or Alu mobilization. To determine whether our approach captured genes with transposon regulatory potential, we assessed whether our list of greenlisted SNVs was enriched for 1) genes with known TE regulatory capabilities and 2) genes in broader pathways involved in TE regulation (**Figure 2A**). Interestingly, our greenlisted SNVs were significantly (FDR = 3.59E-3) enriched in regions near known L1 expression regulators compared to the background list of all SNVs (**Figure 2B, 2C, 2E**). A few examples of these associations included rs1350516110 (p = 9.35E-12, odds ratio = 0.33) which was upstream of *RHOT1*, rs201619112 (p = 6.24E-9, odds ratio = 0.33) which was downstream of *XPR1*, and rs71475866 (p = 1.65E-8, odds ratio = 2.54) which was upstream of *PFKP*. Overall, we identified 24 greenlisted SNVs that were proximal to 10 genes previously annotated as capable of regulating L1 expression. Our greenlisted SNVs also captured genes involved in regulating L1 transposition, though there was no significant enrichment (FDR = 7.78E-1) (**Figure 2B, 2D, 2E**). A few examples of these associations included rs75237296 (p = 2.14E-9, odds ratio = 0.42) in the *PABPC1* 3’UTR, rs1288384419 (p = 3.17E-8, odds ratio = 0.44) in an intron of *RAD51B,* and rs1471205623 (p = 1.34E-7, odds ratio = 0.42) upstream of *MPHOSPH8*. Here, we identified 5 greenlisted SNVs near 3 genes capable of regulating L1 transposition. Finally, we checked the abundances of SNVs linked to candidate regulators of L1 RNA levels in lymphoblastoid cells [41]. However, we were not able to detect any greenlisted SNVs in regions containing candidate genes (**Figure 2B** and **2E**). Nevertheless, these results highlight the ability of our approach to enrich for genomic regions containing known regulators of the retrotransposon lifecycle and suggest that these regulators may play important roles in diverse human populations.

**Fig 2.**
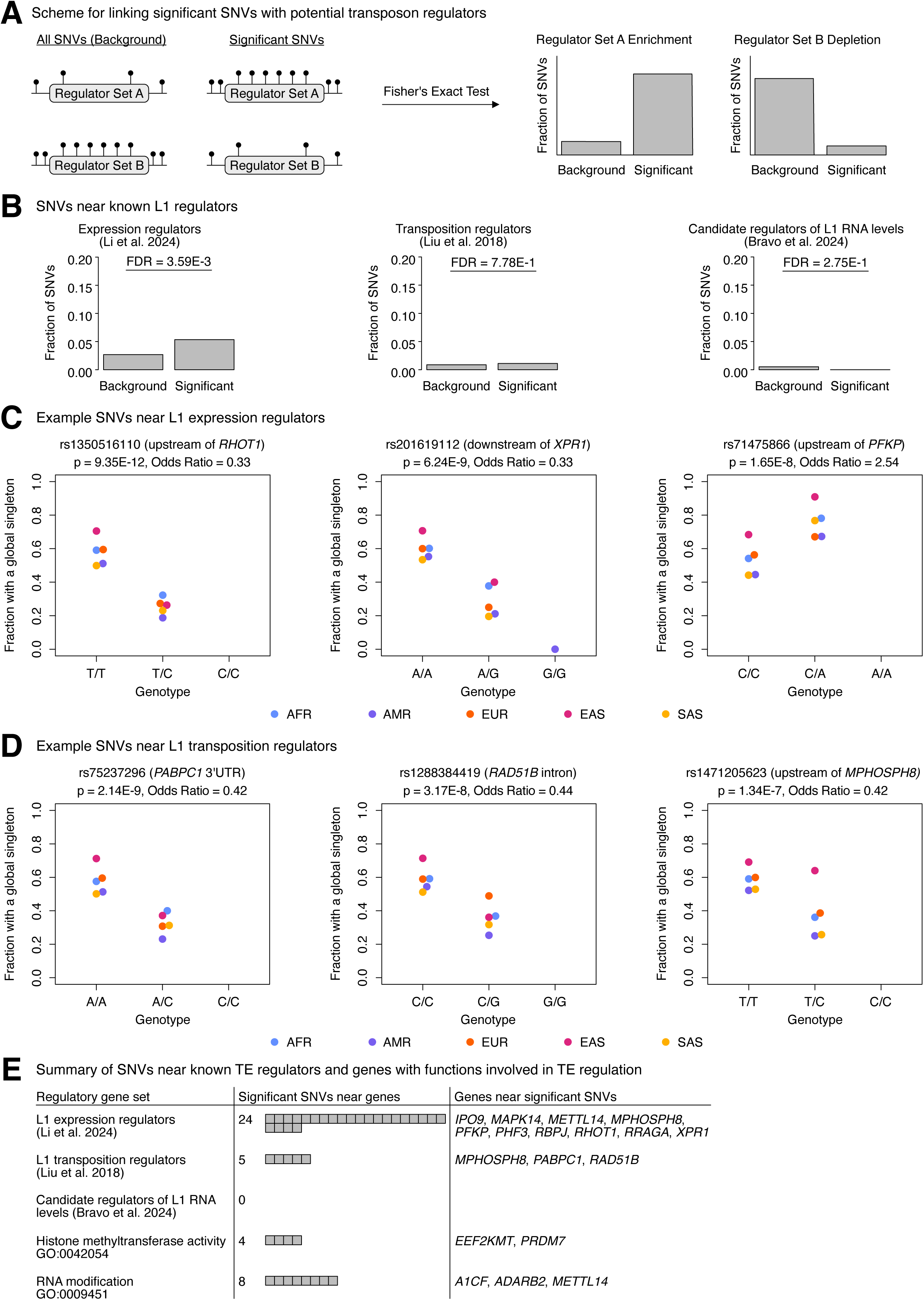
Significant SNVs lie in genomic regions containing genes involved in transposon control. **(A)** Scheme for assessing whether greenlisted SNVs were enriched in regions containing genes with TE regulatory potential. For a given gene set with regulatory potential (regulatory set A or B), the proportion of SNVs near genes in that gene set were calculated for the background and significant SNV lists, and statistical significance was assessed using Fisher’s exact test. **(B)** Enrichment analysis of greenlisted SNVs near genes previously implicated in L1 expression control [37] or L1 transposition control [33] by CRISPR screening in cancer cell lines. Three specific examples of greenlisted SNVs near **(C)** genes controlling L1 expression and **(D)** genes controlling L1 transposition are shown. All GWAS associations have an FDR < 0.05, but, as customary in the GWAS field, we report the raw p-values. **(E)** A summary of the associations we identified with various TE regulatory gene sets, highlighting the number of associated SNVs and the regulatory genes those SNVs were proximal to. Though not exclusive regulators of TE activity *per se*, we included in our analysis gene sets involved in “histone methyltransferase activity” and “RNA modification” functions, since those processes have been implicated in transposon control. FDR: False Discovery Rate.

We next repeated the above analyses using gene sets for broader pathways involved in TE regulation, including a gene set for “histone methyltransferase activity” (GO:0042054) and one for “RNA modifications” (GO:0009451). Though neither gene set was significantly enriched among our greenlisted SNVs (FDR = 1 for methyltransferase activity and FDR = 0.27 for RNA modification), there was some degree of overlap with each gene set (**Figure 2E**). We identified 4 greenlisted SNVs that were proximal to 2 genes with histone methyltransferase activity, including *EEF2KMT* and *PRDM7*. We also identified 8 greenlisted SNVs that were proximal to 3 genes with RNA modification capabilities, including *A1CF*, *ADARB2*, and *METTL14*. Importantly, ADARs (RNA-specific adenosine deaminases) are a family of double-stranded RNA (dsRNA)-binding proteins that modulate A-to-I editing events, including among Alu RNA species, which is important for preventing aberrant activation of innate immune signaling pathways [60]. Though *ADARB2* cannot catalyze A-to-I editing, it can negatively regulate the editing functions of other ADARs [60], making it a potential candidate regulator of Alu activity.

These results demonstrate that our approach can capture genes implicated, but with uncharacterized roles, in TE regulation.

### 2.3 Significant SNVs are enriched in regions containing features that promote genome instability

After scanning for known and potential regulators of TE activity, we next explored the possibility that significant variants tagged genetically unstable TE loci (**Figure 3A**). Such loci could theoretically contribute to TE insertion number variation through *de novo* transposition events. To probe this possibility, we assessed whether greenlisted SNVs were enriched for Alu and L1 loci belonging to subfamilies that have retained their ability to mobilize (**Figure 3B**). Interestingly, SNVs overlapping the active AluS subfamily were significantly (FDR = 4.80E-6) enriched in our SNV greenlist compared to background (**Figure 3B**). Our enrichment analysis also highlighted a significant (FDR = 7.29E-7) depletion of inactive-L1M-overlapping SNVs and a significant (FDR = 3.46E-12) enrichment of L1PA-overlapping SNVs, all compared to background SNVs (**Figure 3B**). To obtain a higher resolution view of the transposition capabilities of overlapping L1PA copies, we checked the overlap of our greenlisted SNVs with annotations for putatively active L1 copies. Surprisingly, active copies, with either an intact, full-length L1 or only an intact ORF2, were not significantly enriched/depleted in our greenlisted SNV list (**Figure 3C**). However, non-intact, full-length L1 copies—annotated for their regulatory potential—were significantly (FDR = 2.41E-6) enriched compared to background. These results suggest that most greenlisted SNV-overlapping L1 copies are limited in their ability to generate *de novo* insertions and thus may influence insertion number through alternative mechanisms (i.e. genome remodeling). This is potentially in contrast to greenlisted SNV-overlapping AluS copies, which may still be measurably active in the human genome. The TE enrichments we identified above were consistent with those identified using the Transposable Element Enrichment Analyzer (TEENA) tool [61] (**Supplementary Table S1H**).

**Fig 3.**
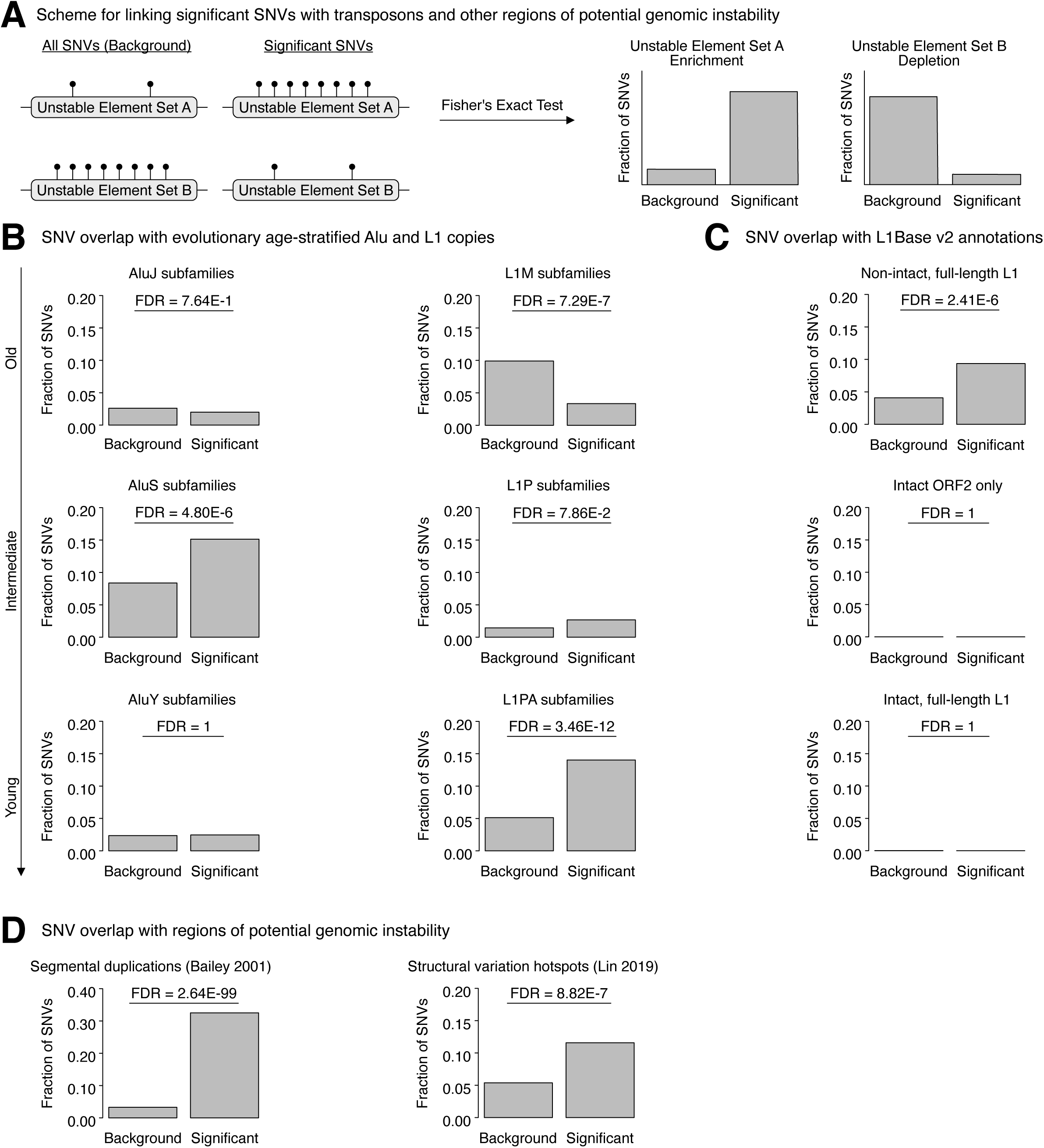
Significant SNVs are enriched in genomic regions containing features associated with genome instability. **(A)** Scheme for assessing whether greenlisted SNVs are enriched in regions containing elements known for promoting genome instability. For a given set of potentially genetically unstable regions (unstable element set A or B), the proportion of SNVs overlapping regions in each set are calculated for the background and significant SNV lists, and statistical significance is assessed using Fisher’s exact test. **(B)** Enrichment analysis of greenlisted SNVs overlapping evolutionary age-stratified Alu (*left column*) and L1 (*right column*) copies. **(C)** Enrichment analysis of greenlisted SNVs overlapping curated L1 loci in L1Base v2 [107]. This database contains putatively active L1 copies (with either full-length, fully intact L1 copies or L1 copies with only ORF2 intact), as well as non-autonomous, full-length, non-intact L1 copies with regulatory potential. **(D)** Enrichment analysis of greenlisted SNVs overlapping genomic regions containing segmental duplications [66, 67] or structural variation hotspots [65]. FDR: False Discovery Rate.

We further explored the more general possibility that greenlisted SNVs tagged genomic regions that may be prone to genome remodeling that may influence TE insertion numbers. Such structural alterations may be facilitated by, but may not require, the presence of repetitive elements. In particular, extensive homology between segmental duplications, often in the vicinity of Alu elements [62], can facilitate NAHR and drive recurrent genomic rearrangements [63] that can help form SV hotspots [64, 65]. Noting the enrichment of AluS copies we observed among our greenlisted SNVs, we next assessed whether our greenlisted SNVs significantly overlapped regions of segmental duplication [66, 67] or regions characterized as SV hotspots [65] (**Figure 3D**). Consistent with the notion that SNVs may tag regions with potentially elevated rates of genome instability, our SNVs were very significantly enriched in regions of segmental duplication (FDR = 2.64E-99), as well as in regions harboring SV hotspots (FDR = 8.82E-7). These results further link variation in TE insertion numbers to genomic loci where structural instability may arise through genome remodeling mechanisms.

Finally, we note that our GWAS analysis identified 21 polymorphic SVs that were significantly (FDR < 0.05) associated with the presence/absence of L1/Alu insertion singletons. These polymorphic SVs varied in nature and included inversions, Alu insertions, an L1 insertion, SINE-VNTR-Alu (SVA) insertions, and a multiallelic copy number variant (**Figure S5A**). With the exception of the CN0 copy number variant (YL_CN_PEL_1784, p = 3.30E-6, odds ratio = 0.28) which was associated with lower odds of carrying an L1/Alu insertion singleton, all of the other structural variants were associated with higher odds of carrying an insertion singleton (odds ratio > 1). Since the sequences for these SVs were not available, it is unclear whether common, polymorphic L1/Alu insertion SVs may be directly increasing the singleton number through novel transposition events, or whether any of these polymorphic SVs may be influencing the L1/Alu singleton number through genome remodeling mechanisms. Indeed, polymorphic inversions, many of which are often flanked by retrotransposons, are associated with genetic instability [68]. Ultimately, these results suggest a tight link between common, polymorphic SVs of different types and L1/Alu singleton SVs, whereby having the former is generally associated with higher odds of having the latter.

### 2.4 Case samples exhibit elevated cell cycle-related gene expression profiles

To gain insight into the functional differences between controls and cases, we leveraged publicly available lymphoblastoid cell line mRNA-seq data generated by the GEUVADIS consortium for a subset of European and African samples in the 1000 Genomes Project [69] (**Figure 4A**). This included 358 Europeans samples from 4 populations (British, Finnish, Tuscan, and Utah residents with European ancestry) and 86 African samples from 1 population (Yoruba), which we recently used to quantify gene and TE expression profiles [41]. We utilized this expression data to construct consensus gene co-expression networks for both the European and African samples using the WGCNA [70] package. This approach led to the identification of 20 consensus modules and 1 module (MEgrey) containing genes that were not assigned to the consensus modules (**Supplementary Table S1I**). We then ran a module-trait correlation analysis comparing the expression of these modules with the case/control status of the European and African samples (**Figure 4B**). Here, we used a stricter threshold of p < 0.01 to call significant correlations. We were not able to identify any significant module-phenotype correlations using the European network, which is potentially consistent with our difficulty in calling significant GWAS variants in this super-population at the available sample sizes (**Figure S2C**). In contrast, the MEroyalblue module was significantly (p = 4.0E-4) correlated with African case/control status. To combine the results from each network, we utilized Fisher’s method to meta-analyze the p-values for modules exhibiting correlations in the same direction. By meta-analysis, the MEroyalblue module was still significantly (p = 0.002) and positively correlated with case status. Finally, to functionally characterize this module, we ran over-representation analysis using the GO Biological Process gene set collection (**Figure 4C, Supplementary Table S1J**). The top 10 over-represented gene sets were involved in cell cycle-related processes, including “mitotic cell cycle”, “cell division”, and “sister chromatid segregation”. These findings are consistent with the biology underlying an expansion of TE insertions. Though L1 can mobilize in non-dividing cells [71, 72], L1 retrotransposition exhibits a cell cycle bias and peaks during the S phase [73]. Alternatively, chromosome segregation errors during mitosis or meiosis can generate cells with abnormal ploidy and either increased or decreased dosages of both genic and transposon content [74]. These results implicate cell cycle differences in cells from individuals with unique L1/Alu insertion singleton variation.

**Fig 4.**
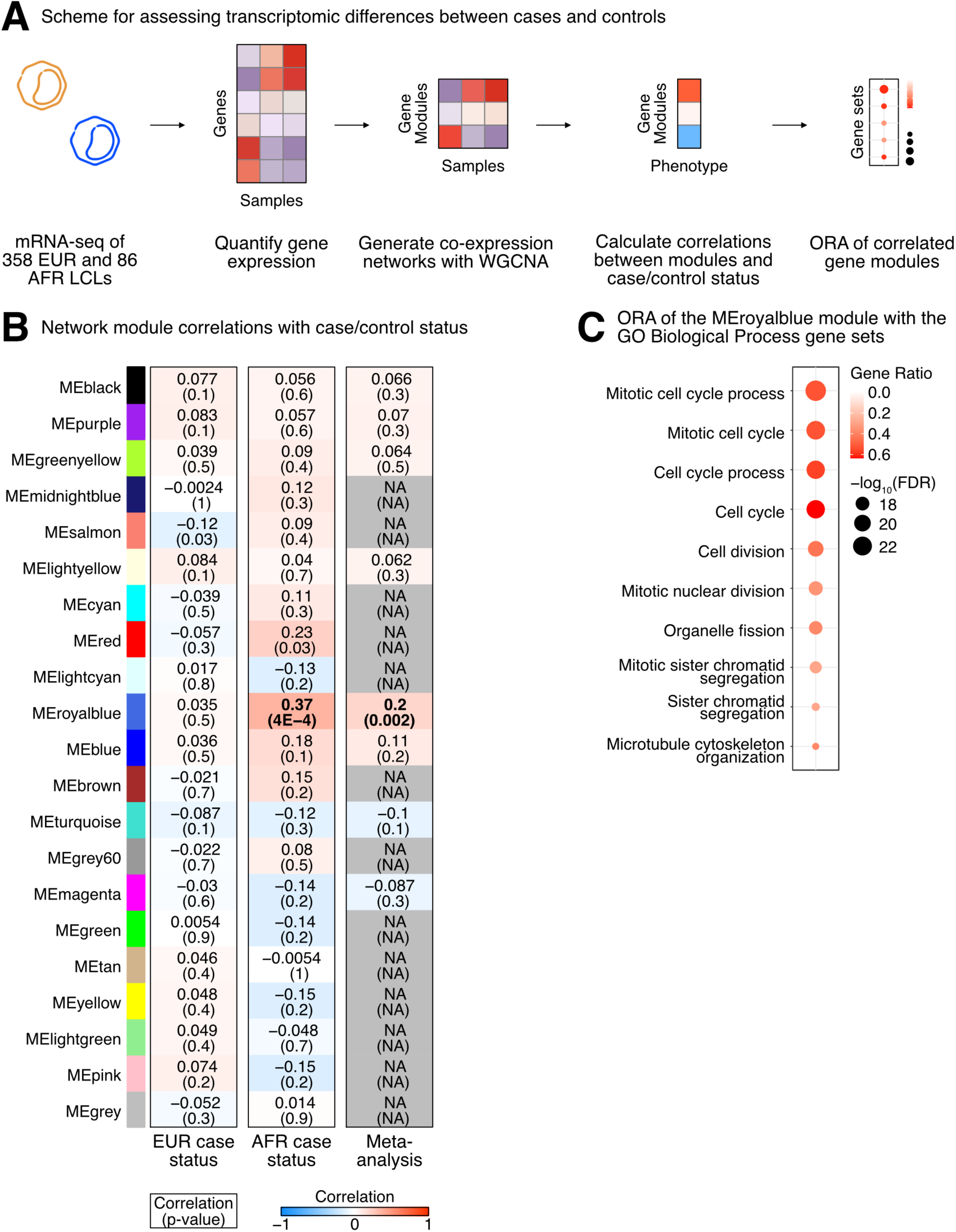
Alterations in the cell cycle are positively correlated with case status. **(A)** Scheme for characterizing transcriptomic differences between case and control samples. Gene expression profiles were quantified using mRNA-sequencing data from lymphoblastoid cells belonging to 358 European and 86 African individuals. To note, all African individuals here were from the Yoruban population. These gene expression profiles were used to construct consensus gene co-expression networks with WGCNA. We then quantified the correlations between each module in the network and the case-control status of all samples (encoded as 0 for controls and 1 for cases). Finally, over-representation analysis (ORA) using the Gene Ontology (GO) Biological Process gene set collection was used to assign functions to significantly correlated modules. **(B)** The correlations and correlation p-values between consensus network modules and case-control status in the European and African cohorts. Boxes were color-coded according to the strength of the correlation. A meta-analysis was also carried out to summarize statistical results by combining European and African correlation p-values using Fisher’s method. For visualization purposes only, the average of the European and African correlations was assigned to the meta-analysis. Correlations with opposite trends in the two cohorts were disregarded in the meta-analysis and colored grey. Correlations with p < 0.01 were considered statistically significant and highlighted in bold. **(C)** The top 10 ORA results for the MEroyalblue module using the GO Biological Process gene set collection. The colors represent the gene ratio (i.e. the fraction of module genes from the listed gene set) and the sizes of the dots represent the Benjamini-Hochberg FDR. NA: Not Applicable. FDR: False Discovery Rate.

## 3. DISCUSSION

### 3.1 A new approach to identify loci implicated in L1 and Alu insertion number variation

In this work, we developed a pipeline to computationally identify candidate loci involved in L1/Alu singleton number variation by GWAS analysis. Importantly, our study incorporates natural human genetic variation present in populations of different geographic origin via trans-ethnic GWAS meta-analysis to identify shared, candidate regulatory loci. Though several studies have begun to screen for regulators and potential regulators of L1 expression or transposition in cell culture models or across tissues [33–40], these can be limited in that the generalizability of these findings to different ethnic populations is unclear. Moreover, no systematic, genome-wide screen for candidate regulators of Alu activity has been carried out thus far, to our knowledge. To address these gaps, we previously utilized trans-eQTL analysis to identify potential regulators of L1 RNA levels in European and African populations [41]. Here, we utilized genomic data from samples originating from 5 super-populations to identify candidate loci modulating L1/Alu insertion singleton numbers.

TE insertion number variation can arise through *de novo* transposition events or through genome remodeling mechanisms that can generate large deletions or duplications. We were particularly interested in identifying new candidate regulators of L1/Alu transposition. Consistent with the notion that greenlisted SNVs may play roles in the retrotransposon lifecycle, our approach enriched genomic regions containing genes that can regulate L1 expression levels. Though other known regulators of TE activity, and pathways involved in TE control, were not significantly enriched among our greenlisted SNVs, we nonetheless identified many SNVs in genomic regions containing these genes. This included, for example, *MPHOSPH8*—a component of the HUSH complex important for L1 repression, regulating both L1 expression and transposition [33, 37, 75–77]. It also included *PABPC1* which is a poly(A) binding protein that attaches to the poly(A) tail of L1, is important for the formation of L1 ribonucleoprotein particles (RNPs), and modules L1 and Alu transposition [78–80]. As a third example, we identified variants near *ADARB2*, a negative regulator of RNA A-to-I editing, including among Alu RNAs [60]. These results suggest that SNV-associated genes identified in this study hold TE regulatory potential and it may therefore be informative to (i) test and validate these in future studies or (ii) use these to prioritize future, targeted studies of TE regulators.

Our approached also identified an enrichment of greenlisted SNVs in regions containing reference TE insertions, including AluS and full-length, non-intact L1PA copies. Though neither of these can mobilize autonomously, they can hijack machinery from transposition-competent L1s and mobilize *in trans* [6, 7, 9]. Thus, it is possible that greenlisted SNVs tag reference insertions contributing to L1/Alu singleton variation through transposition-dependent mechanisms. Of course, an alternative possibility is that these repetitive elements are directly involved in genomic remodeling involving mechanisms like repeat-mediated deletions or NAHR. We also note that greenlisted SNVs were enriched in regions containing segmental duplications and structural variation hotspots where recombination-based mechanisms, including NAHR, may lead to duplications or deletions of the local genomic architecture. Thus, it is also possible that greenlisted SNVs tag genomic regions prone to structural variation that can alter the L1/Alu insertion number through recombination-dependent mechanisms.

Importantly, we also suggest the possibility that genome remodeling mechanisms (including recombination) may interact with gene-based mechanisms of TE regulation. Indeed, genes such as *BRCA1* are known to regulate L1 transposition [33] and are also known to undergo Alu-Alu recombination events that can give rise to new mutations in the gene [81–83]. Observations such as these highlight the possibility that TE insertions may modulate structural variation in genomic regions containing genes regulating retrotransposon lifecycles, which may facilitate an expansion of TE insertions through transposition-based mechanisms, which may influence further structural variation driving this whole process. This possibility is consistent with the enrichment of greenlisted SNVs in regions containing L1 expression regulators, Alu and L1 repeats, and other genomically unstable features like segmental duplications and structural variation hotspots. In the future, it may be informative to experimentally assess this possibility in contexts where genome instability is a hallmark feature that is coupled with TE de-repression, such as aging [21] or aging-associated diseases like cancer [84, 85].

Finally, our approach also identified several common, polymorphic SVs that were significantly associated with L1/Alu insertion singletons. Overwhelmingly, the presence of polymorphic SVs of different types—inversions, Alu insertions, an L1 insertion, and SVA insertions—was associated with increased odds of a global L1/Alu insertion singleton. The exception to this was a multi-allelic copy number variant (YL_CN_PEL_1784, p = 3.30E-6, odds ratio = 0.28) where 0 copies were present; this SV was associated with decreased odds of a global L1/Alu insertion singleton. Based on these results, we speculate that specific polymorphic SVs (i) may directly drive genome instability that can facilitate the acquisition of L1/Alu copies or (ii) may serve as markers for elevated risk of indirectly acquiring additional L1/Alu copies. Indeed, active donor L1 copies that can mobilize and generate *de novo* insertions are usually highly polymorphic in human populations (reviewed in [59]), and polymorphic inversions, many of which are often flanked by retrotransposons, are also associated with genetic instability and genomic disorders [68].

In summation, this study provides a list of variants that are associated with L1/Alu insertion singletons and includes (i) SNVs in regions containing regulators of TE activity, (ii) SNVs in regions containing features associated with genome instability, including retrotransposons, that may influence TE insertion number variation through transposition-dependent or transposition-independent mechanisms, and (iii) common, polymorphic SVs that may also influence TE insertion number variation through transposition-dependent or transposition-independent mechanisms.

### 3.2 L1/Alu insertion singleton-associated loci contain genes of clinical relevance

The most significant, greenlisted variant we identified was a polymorphic inversion SV (INV_delly_INV00066128, chr21:26001780, p = 5.67E-23, odds ratio = 4.38) residing in an intronic Alu copy within the *APP* gene, an important marker of Alzheimer’s disease (AD). AD is characterized by (i) the accumulation of amyloid β (Aβ) plaques derived from amyloidogenic *APP* processing and (ii) neurofibrillary tangles of hyperphosphorylated tau [86]. Importantly, tau protein can induce TE expression and there is speculation that TEs may mobilize in tauopathies [86]. Whether APP protein or Aβ plaques can similarly modulate TE expression or mobilization is an interesting area of potential future research; indeed, our findings are consistent with the possibility that *APP* products may act as regulators of TE activity. Of course, another possibility is that the genomic region containing *APP* may be a source of L1/Alu insertion number variation independent of the functional properties of APP protein (i.e. genomic instability at that locus may be the driver of TE insertion number variation). Indeed, repeat-based recombination events have been documented in AD [87]. Nevertheless, our results offer another connection between transposable element regulation and Alzheimer’s disease.

We also identified several SNVs proximal to *PWRN1*, which resides within the Prader-Willi syndrome region and is thought to play a role in PWS [88]. Prader-Willi is an imprinting disorder where genes in the chromosome 15q11-q13 region are maternally imprinted and paternal copies are not expressed [89]. This lack of paternal gene expression is predominantly caused by *de novo* paternally inherited deletions of the 15q11-q13 region, though, less frequently, inheritance of two maternal chromosome 15 copies is the cause [89]. Importantly, a feature of genomic disorders like PWS is the presence of segmental duplications that can serve as substrates for NAHR [90, 91]. Thus, we hypothesize that this particular region might be more prone to L1/Alu insertion number variation as a consequence of recombination-based chromosomal alterations. Nevertheless, it is unclear (i) whether L1/Alu repeats are differentially active in PWS compared to healthy controls or, more specifically, (ii) whether genes like *PWRN1* can differentially express or mobilize L1/Alu transposons. Ultimately, the associations between L1/Alu singletons and both *APP* and *PWRN1* further implicate retrotransposons in disease.

Though we did not identify aging-specific signatures in our analyses, these results should nonetheless be relevant for carrying out a comprehensive analysis of aging-associated genetic risk factors. Though further validation work is needed, it is hypothesized that L1 insertions increase with chronological aging [22] and during cellular senescence [23]. Moreover, both L1 and Alu can promote features of cellular senescence [26–30]. Though these associations are increasingly being characterized, the study of transposable elements has generally been limited, and their relationship with aging processes remain incompletely characterized. Thus, this study may serve to accelerate identification of these relationships by providing an initial set of high-confidence variants that can be incorporated into future genetic scans for aging-associated risk factors. Given the ever-increasing availability of large-cohort-based association studies, these analyses should provide immediate, human-relevant, and novel aging molecular targets.

### 3.3 Limitations and future considerations

In this study, we sought to identify new, candidate genes implicated in Alu and L1 insertion number control. One specific mechanism of interest by which this can occur is target-primed reverse transcription (TPRT)-mediated transposition. Since insertion sequences for 1000 Genomes Project samples were not available, to our knowledge, it is difficult to assess to what degree TPRT is driving the associations we identified, since we cannot check for sequence features of TPRT. In studies with larger cohorts where insertion sequences are available and insertions with TPRT features can be identified, our approach could theoretically be applied to explore the genetic basis of TPRT-specific insertion number variation. Of course, our approach is generally restricted to identifying genomic loci where variation is common across human populations. We note, however, that significant variants were enriched in regions containing genes involved in L1 expression regulation. Since expression is one of the early steps of the L1 life cycle, our approach captured variants and genes with potential significance to TPRT-mediated transposition.

We also note that there are several variables that we are unable to control for in this study. To protect patient privacy, biological covariates such as chronological age were not available and therefore could not be corrected for in our analysis. Since increases in L1 expression and L1 copies have been observed with chronological aging [22], differences in insertion number may reflect age differences rather than genetic differences. Importantly, however, samples were considered healthy at the time of sample collection, potentially mitigating health-related effects on insertion number. The origin and developmental timing of the rare Alu and L1 insertions we utilized is also unclear. That is, it is unclear whether global singletons used in this study arose *de novo* in the germline, arose during life through somatic mutation, or even whether they just arose during the cultivation of the lymphoblastoid cells used to amplify each sample’s DNA. Depending on when these insertions were acquired, the associations identified in this study may be relevant for either germ cell or somatic cell TE biology. A potential avenue of future research to address this question would be the incorporation of trio parent-child genome sequencing [92] and multi-generational genome sequencing to help identify *bona fide de novo* insertions and their developmental timing. Additionally, to further strengthen reliability and generalizability of our findings, it will be important to assess whether significant variants identified in this study are also identified in other independent cohorts, such as those in the UK Biobank.

More broadly, though we employed GWAS to scan most of the genome for variants associated with differences in genomic TE content, we note that the X and Y chromosomes were omitted from our scan for several reasons. These included: (1) the necessary data was of poor quality, with over 80% of the X chromosome not genotyped by the 1000 Genomes Project, (2) the necessary data from the Y chromosome was not available, to our knowledge, and (3) the analyses require additional analytical considerations (i.e. male hemizygosity and X chromosome inactivation) that warrant their own study design [93–96]. Though the data availability and data quality limitations have been partially addressed by additional sequencing and variant calling in 1000 Genomes Project samples [92], incorporation of this data and of additional analytical approaches would be beyond the scope of this paper. Thus, the contributions of the X and Y chromosomes to differences in TE genomic load are a ripe area for future investigation.

Ultimately, future studies modulating genes identified with our approach will need to be carried out to validate any causal contributions of the identified variants to L1/Alu genomic load. This validation may take on multiple forms, depending on the hypothesis being tested. For example, if variants tag genes regulating TE mobilization, the tagged genes can be knocked-down or overexpressed and TE mobilization can be assessed *in vitro* using a retrotransposition assay [97]. Alternatively, if the variants tag genomic regions prone to structural remodeling (i.e. contribute to NAHR or segmental duplications), these regions can be sequenced prior to and after a challenge, to assess whether the number of TE copies has been altered. The underlying mechanisms driving differences in genomic TE load will likely be multifactorial.

### 3.4 Conclusions

In this study, we employed GWAS across human populations of different geographic origin to computationally identify genomic loci associated with variation in L1/Alu insertion numbers. Our approach enriches for SNVs in genomic regions containing known regulators of L1 expression. This observation suggests that the TE-regulatory properties of these genes may extend beyond isogenic cell culture models to more diverse human populations. Moreover, this observation also suggests that our list of associated genes likely contains novel regulators of L1 or Alu activity that may be prioritized in future, validation studies. Our approach also identified reference insertions and non-reference, polymorphic SVs that may modulate L1/Alu insertion numbers through transposition-dependent or transposition-independent mechanisms. Finally, the observation that some significant variants reside in genes of clinical relevance, like *APP* and *PWRN1*, reinforce accumulating evidence of biological associations between TE regulation and disease. Ultimately, our approach adds to the analytical toolkit that can be used to study the regulation of TE activity.

## 4. METHODS

### 4.1 Publicly available genomic data acquisition

The multi-ethnic GWAS analysis was carried out on 2503 individuals derived from 5 super-populations (African, East Asian, European, South Asian, and American) and for which paired single nucleotide variant and structural variant data were available from Phase 3 of the 1000 Genomes Project [98–100]. Specifically, Phase 3 autosomal SNVs called on the GRCh38 reference genome were obtained from The International Genome Sample Resource (IGSR) FTP site (http://ftp.1000genomes.ebi.ac.uk/vol1/ftp/data_collections/1000_genomes_project/release/20190312_biallelic_SNV_and_INDEL/). Structural variants, called against the GRCh37 reference genome and then lifted over to GRCh38, were also obtained from the IGSR FTP site (http://ftp.1000genomes.ebi.ac.uk/vol1/ftp/phase3/integrated_sv_map/supporting/GRCh38_positions/).

Human gene expression data across 54 non-diseased tissue sites was obtained from the GTEx Analysis v8 on the GTEx Portal [56–58]. Specifically, we downloaded the matrix containing the median gene-level transcripts per million (TPMs) by tissue, and we extracted the expression values for significant SNV-associated genes. After filtering out genes with no detectable expression (0 TPMs), we generated heatmaps using the pheatmap v1.0.12 package in R. Gene expression values were centered and scaled across tissues to visualize and compare the relative expression levels across tissues.

### 4.2 Aggregating and pre-processing genotype data for GWAS analysis

To define the phenotype of interest for GWAS, we first extracted global singleton Alu and L1 insertions. We utilized VCFtools v0.1.17 [101] to extract all autosomal SVs with no missing data (--max-missing 1) and an allele count of 1 across all samples (-- non-ref-ac 1 --max-non-ref-ac 1), i.e. global singletons. From these, we extracted Alu- and L1-specific insertions using BCFtools v1.10.2 [102] to keep entries annotated with the ‘SVTYPE=“LINE1”’ and ‘SVTYPE=“ALU”’ tags. We calculated the enrichment of various genomic features overlapping L1/Alu singletons using the annotatePeaks script within the Hypergeometric Optimization of Motif EnRichment (HOMER) v4.11.1 software suite [103] using hg38 v6.4 annotations. Finally, VCF files containing global singleton L1 or Alu insertions were converted to PLINK BED format using PLINK v1.90b6.17 [104]. We note that SVs on sex chromosomes were not included in any part of the analysis since (1) the necessary data was of poor quality, with over 80% of the X chromosome not genotyped by the 1000 Genomes Project, (2) the necessary data from the Y chromosome was not available, to our knowledge, and (3) the analyses require additional analytical considerations (i.e. male hemizygosity and X chromosome inactivation) that warrant their own study design [93–96].

Secondly, we prepared polymorphic SVs for inclusion in the GWAS analysis. VCFtools was used to isolate SVs with the following properties in each individual super-population: possessed a minimum and maximum of two alleles (biallelic), possessed a minor allele frequency (MAF) of at least 1%, passed Hardy-Weinberg equilibrium thresholding at p < 1e-6, had no missing samples, and was located on an autosome. To focus on shared, trans-ethnic sources of genetic variation, we used BCFtools to identify and subset SVs that were shared across all 5 super-populations.

Third, we prepared polymorphic SNVs for inclusion in the GWAS analysis. All SNVs were first annotated with rsIDs from dbSNP build 155 using BCFtools. Within each super-population, VCFtools was used to remove indels and keep autosomal SNVs with the same parameters as the polymorphic SVs. We note that for similar reasons as with the polymorphic SVs, sex chromosome SNVs were also omitted from all analyses. We then used BCFtools to identify and subset SNVs that were shared across all 5 super-populations. Finally, we used BCFtools to generate the final genotype matrices by combining shared, polymorphic SNVs with shared, polymorphic SVs. VCF files containing the combined SNVs and SVs were then converted to PLINK BED format using PLINK, keeping the allele order. PLINK was also used to prune the combined SNV and SV matrices (--indep-pairwise 50 10 0.1) and to generate principal components (PCs) from the pruned genotypes, for inclusion as covariates in the GWAS.

### 4.3 Super-population-specific and trans-ethnic GWAS

We began by running GWAS within each super-population using PLINK v1.90b6.17 [104]. For the phenotype, we added the number of Alu and L1 global singleton insertions for each sample and segregated samples into cases and controls, depending on whether they contained or did not contain a global singleton. We ran GWAS analyses using a logistic regression model that included the following covariates: biological sex and the top 4 principal components from the pruned SNV and SV genotype matrices. Individual results from each super-population were combined via meta-analysis using PLINK. To help call significant variants, we generated a null distribution of p-values for each super-population by running 20 instances of the GWAS where the case-control status for each sample was randomly shuffled with the case-control status of a different sample. Each set of permutation results was meta-analyzed across super-populations to similarly obtain 20 random distributions of meta-analysis p-values. For the meta-analysis, we focused on the p-values and odds ratios generated using a random effects statistical model, as opposed to a fixed effects model, since 1) there may be heterogeneity across super-populations in response to different genetic variants, and 2) we were interested in enhancing the generalizability of our findings to facilitate future follow-up studies.

To limit false positives, the Benjamini-Hochberg (BH) false discovery rate (FDR) was calculated in each analysis, and we used the p-value corresponding to a BH FDR < 0.05 as the threshold for GWAS significance. As a secondary threshold, we used the permutation data to identify p-values corresponding to an average empirical FDR < 0.05. To note, we calculated the average empirical FDR at a given p-value p_i_ by (i) counting the total number of null points with p ≤ p_i_, (ii) dividing by the number of permutations, to obtain an average number of null points with p ≤ p_i_, and (iii) dividing the average number of null points with p ≤ p_i_ by the number of real points with p ≤ p_i_. GWAS variants were considered significant if they passed the stricter of the two thresholds in each analysis.

### 4.4 Annotation of variants and annotation enrichment analyses

We obtained BED files containing annotated genomic regions from various sources. We obtained the ENCODE blacklist v2 [52] for hg38 from https://github.com/Boyle-Lab/Blacklist/tree/master/lists. This curated “blacklist” represents a set of genomic regions with anomalous signal in next-generation sequencing experiments independent of cell type and individual experiment [52]. SNVs overlapping these regions were significantly enriched in our initial, unfiltered significant SNV list compared to the background SNV list (Fisher’s exact test, FDR = 1.33E-302). Segmental duplications [66, 67] and RepeatMasker annotations using the Repbase library [105] were obtained from the UCSC Genome Browser [106]. We obtained SV hotspot coordinates on hg19 from [65], and we used the online UCSC LiftOver tool to map coordinates to the hg38 genome assembly using the default settings. The BED tracks for full-length and intact L1s, only ORF2-intact L1s, and full-length non-intact L1s were obtained from L1Base v2—a dedicated database of putatively active L1 insertions [107]. We obtained the Registry (version 4) of candidate cis-Regulatory Elements (cCREs) for hg38 from the Search Candidate Regulatory Elements by ENCODE (SCREEN) web interface [55] (http://screen-beta.wenglab.org). We used the ‘intersect’ command in BEDTools v2.31.1 [108] to assign genomic region annotations to all overlapping variants used in this study. We were also interested in assessing whether variants were linked to specific regulatory gene annotations. All variants used in the study were submitted to the GREAT v4.0.4 [109, 110] online platform with the default settings (basal plus extension, proximal with 5 kb upstream and 1 kb downstream, plus distal up to 1000 kb) to assign gene annotations to each variant. These gene annotations were then used to assess the number of variants linked to genes in several TE regulatory lists—including a list of genes that regulated L1 expression in a CRISPR screen using cancer cells [37], a list of genes that regulated L1 transposition in an independent CRISPR screen using cancer cells [33], a list of candidate genes influencing intronic, intergenic, or exonic L1 RNA levels in lymphoblastoid cell lines [41], a list of genes with histone methyltransferase activity (GO:0042054), and a list of genes with RNA modification activity (GO:0009451). The two GO gene sets were obtained on 2024-07-29 from the Molecular Signatures Database (MSigDB) v2023.2.Hs [111, 112], corresponding to GO release 2023-07-27.

Given the limited number of significant SVs and the unavailability of SV sequences, we largely focused on blacklist-filtered, significant SNVs in downstream enrichment analyses. For each of the above annotations, we assessed whether greenlisted SNVs were significantly enriched or depleted for that annotation compared to background SNVs—all SNVs that were tested in the GWAS. The numbers of background and greenlisted SNVs overlapping or not overlapping a set of annotations were placed into a contingency table, and statistical significance was assessed using Fisher’s exact test (with the fisher.test function in R v4.3.3). After all tests were carried out, p-values were FDR corrected using the p.adjust function in R. All enrichments/depletions with an FDR < 0.05 were considered significant. For the repeat enrichment analyses, we also analyzed our greenlisted SNVs using the TEENA web server [61] (on August 8, 2024), specifying the hg38 assembly and using all other default options.

### 4.5 RNA-seq and gene co-expression network analyses

For lymphoblastoid cell line transcriptional analyses, mRNA-sequencing was initially carried out by the GEUVADIS consortium [69] on LCLs from a small subset of European and African (Yoruban, specifically) samples from the 1000 Genomes Project. Recently, we re-processed this data to quantify gene and transposon expression levels [41]. Briefly, reads were trimmed using fastp v0.20.1 [113], trimmed reads were aligned to the GRCh38 human genome assembly using STAR v2.7.3a [114], and the TEtranscripts v2.1.4 [115] package was used to obtain gene and TE counts, using the GENCODE release 33 [116] annotations and a repeat GTF file provided on the Hammell lab website. To note, the EBV genome (GenBank ID V01555.2) was included as an additional contig in our reference genome, since LCLs are generated by infecting B-cells with Epstein-Barr virus (EBV).

Using these gene/TE count matrices, lowly expressed genes were filtered out if 50% of European or Yoruban samples did not have over 0.44 counts per million (cpm) or 0.43 cpm, respectively, which correspond to 10 reads in each cohort’s median-length library. Since we were interested in building consensus co-expression networks between the European and Yoruban samples, we also removed genes that were not expressed in both groups. After, the filtered counts underwent a variance stabilizing transformation (vst) using DESeq2 v1.42.1 [117] and the following covariates were regressed out using the ‘removeBatchEffect’ function in Limma v3.58.1 [118]: biological sex, sequencing lab, population category, principal components 1-2 of the pruned genotype matrices containing both SNVs and SVs, and EBV expression levels. The population category variable was omitted in the Yoruban batch correction since that did not vary.

The batch-corrected VST matrices were then used to perform weighted gene co-expression network analysis (WGCNA) [70] using the WGCNA v1.72-5 R package. We used the ‘blockwiseConsensusModules’ function to automate consensus network construction for both the European and African expression data, specifying these parameters: corType = “bicor”, power = 12, networkType = "signed", maxPOutliers = 0.05, mergeCutHeight = 0.25, deepSplit = 2, minKMEtoStay = 0, pamRespectsDendro = FALSE, minModuleSize = 30, and randomSeed = 90280. Phenotype-module correlations, and the corresponding p-values, were calculated using WGCNA’s ‘cor’ and ‘corPvalueFisher’ functions, respectively. The p-values for the European and Yoruban correlations were meta-analyzed using Fisher’s method. For visualization purposes only, to show a correlation direction in the meta-analysis, we took the average of the European and Yoruban correlations. Modules showing opposite correlations across the two consensus networks were disregarded in the meta-analysis. Correlations with a p-value < 0.01 were considered significant.

### 4.6 Functional enrichment analyses

We used the over-representation analysis (ORA) paradigm as implemented in the R package clusterProfiler v4.10.1 [119]. Gene Ontology Biological Process gene sets were obtained from the R package msigdbr v7.5.1, an Ensembl ID-mapped collection of gene sets from the Molecular Signatures Database [111, 112]. For ORA with genes linked to greenlisted SNVs and SVs, we used the genes linked to the background SNVs and SVs, respectively, for the universe background to compute enrichment significance. For ORA analysis of co-expression network modules, we used all genes in the network for the universe background. All gene sets with an FDR < 0.05 were considered significant, and the top 10 significant gene sets, at most, were plotted. All enrichments results are included in **Supplementary Table S1D, S1E,** and **S1J**.

## DECLARATIONS

### Ethics approval and consent to participate

Not applicable.

### Consent for publication

Not applicable.

### Availability of data and materials

All code is available on the Benayoun lab GitHub (https://github.com/BenayounLaboratory/TE_GWAS). Analyses were conducted using R version 4.3.3. Code was re-run independently on R version 4.3.0 to check for reproducibility.

### Authors’ contributions

**Juan I Bravo:** Conceptualization, formal analysis, investigation, methodology, visualization, writing - original draft preparation, writing - review and editing. **Lucia Zhang:** Validation, formal analysis, writing - review and editing. **Bérénice A. Benayoun:** Conceptualization, formal analysis, funding acquisition, methodology, supervision, visualization, writing - original draft preparation, writing - review and editing.

## Acknowledgements

We would like to thank Prof. Rachel Brem at the University of California Berkeley for her feedback and insights on the GWAS analyses. We would also like to thank Dr. Heather C. Mefford at St. Jude Children’s Research Hospital for referring us to publications that helped shape the analyses linking our variants to regions potentially prone to genomic instability and involved in disease. Finally, we would like to thank Dr. Minhoo Kim, Mr. Aaron Lemus, and Ms. Eyael Habtemariam for their comments on the manuscript.

## Legends to Supplementary Figures

**S1 Fig. Quality control of combined SNV and SV 1000 Genomes Project data.**

PCA plots for pruned SNV and SV genotype data from **(A)** African, **(B)** East Asian, **(C)** European, **(D)** South Asian, and **(E)** Admixed American samples. Colors and shapes represent different populations within each super-population.

**S2 Fig. GWAS in individual super-populations is underpowered.**

Manhattan plots for GWAS results in individual super-populations, including for the **(A)** African, **(B)** East Asian, **(C)** European, **(D)** South Asian, and **(E)** Admixed American cohorts. Solid lines correspond to a Benjamini-Hochberg FDR < 0.05 and dashed lines correspond to an average empirical FDR < 0.05, based on 20 random permutations. The Benjamini-Hochberg FDR and average empirical FDR, respectively, corresponded to the following p-values in each super-population: p = 1.18E-6 and p = 4.61E-6 in the African cohort, p = 9.06E-7 and p = 8.46E-7 in the South Asian cohort, and p = 3.53E-7 and p = 1.07E-6 in the Admixed American cohort. The stricter of the two thresholds in each super-population was used to define significant SNVs and SVs. No variant at an FDR < 0.05 was identified in the East Asian and European cohorts. Significant variants overlapping regions in the ENCODE blacklist v2 are shown in blue. FDR: False Discovery Rate.

**S3 Fig. GWAS meta-analysis associations are conserved in non-African super-populations.**

**(B)** A schematic illustrating the trans-ethnic integration of available SNV and SV data to identify variants associated with L1/Alu insertion global singletons in non-African super-populations. Within each non-African super-population, samples were segregated into cases and controls depending on whether or not they harbored a global Alu or L1 insertion singleton. GWAS was carried out within each super-population to identify polymorphic SNVs and SVs associated with case-control status. Finally, GWAS results from all 4 non-African super-populations were meta-analyzed using a random effects statistical model, yielding a summary meta-analysis odds ratio and p-value for each variant. **(B)** A Manhattan plot for the trans-ethnic GWAS meta-analysis omitting the African super-population. The dashed line at p = 6.60E-6 corresponds to an average empirical FDR < 0.05, based on 20 random permutations. One such permutation is shown in the bottom panel for illustrative purposes. The solid line at p = 1.92E-6 corresponds to a Benjamini-Hochberg FDR < 0.05. The stricter of the two thresholds, p = 1.92E-6, was used to define significant SNVs and SVs. Significant variants overlapping regions in the ENCODE blacklist v2 are shown in blue and were omitted from downstream analyses. **(C)** A Venn diagram comparing the number of significant variants identified using either all five super-populations or all four non-African super-populations. The statistical significance of the overlap was calculated using a one-sided Fisher’s exact test. **(D)** A Venn diagram comparing the number of significant variants identified using either all four non-African super-populations or only the African super-population. The statistical significance of the overlap was calculated using a one-sided Fisher’s exact test. FDR: False Discovery Rate.

**S4 Fig. Functional annotation of significant variants.**

**(A)** Scheme for predicting functions of genomic regions containing significant variants. All SNVs and SVs used in this study were assigned genes using the GREAT [109] online platform. Significant SNV- and SV-associated genes were then tested for functional gene set enrichment by over-representation analysis (ORA) using clusterProfiler [119], specifying the corresponding background-associated genes as the universe. **(B)** The number of genes associated to greenlisted SNVs (*left*), the distance between greenlisted SNVs and the transcriptional start sites (TSS) of associated genes (*middle*), and the top 10 ORA results for associated genes using the GO Biological Process gene set collection (*right*). The colors represent the gene ratio (i.e. the fraction of significant SNV-associated genes from the listed gene set) and the sizes of the dots represent the Benjamini-Hochberg FDR. **(C)** The number of genes associated to greenlisted SVs (*left*) and the distance between greenlisted SVs and the transcriptional start sites (TSS) of associated genes (*middle*). **(D)** Enrichment analysis of greenlisted SNVs overlapping genomic regions with candidate cis-Regulatory Elements (cCREs) from the ENCODE Registry v4 [55]. **(E)** Heatmap comparing the median expression levels of significant SNV-associated genes in each tissue included in the GTEx Analysis v8. Values in the heatmap represent z-scores of gene expression across tissues. Negative values are lower than average tissue expression and positive values are higher than average tissue expression. FDR: False Discovery Rate.

**S5 Fig. Polymorphic SVs of different classes are associated with L1/Alu insertion singletons.**

**(A)** One example of each type of significant, polymorphic SV that was associated with L1/Alu singletons. These classes included inversions, Alu insertions, an L1 insertion, SINE-VNTR-Alu (SVA) insertions, and a multiallelic copy number variant. All GWAS associations have an FDR < 0.05, but, as customary in the GWAS field, we report the raw p-values. FDR: False Discovery Rate.

## Inventory of Supplementary Tables

**Supplementary Table S1. Sample singleton counts, significant variant annotations, and gene co-expression network results.**

**(A)** Number of singletons for each sample. **(B)** All variants passing FDR < 0.05 in the GWAS meta-analysis. **(C)** Significant variants uniquely identified in the non-African meta-analysis and absent in the complete meta-analysis. **(D)** Over-representation analysis of genes associated to greenlisted, significant SNVs using GO Biological Process gene sets. **(E)** Over-representation analysis of genes associated to greenlisted, significant SVs using GO Biological Process gene sets. **(F)** SnpEff annotations of significant variants. **(G)** Expression levels (median tissue-specific TPMs) of significant SNV-associated genes in the GTEx Analysis v8. The cluster of brain-associated genes is in blue, and the cluster of testes-associated genes is in orange. **(H)** TE enrichment analysis with TEENA. **(I)** Lymphoblastoid cell line WGCNA network gene-module assignments. **(J)** Over-representation analysis of MEroyalblue genes using GO Biological Process gene sets.

