## Supplementary figures and images for "Multi-ancestry GWAS reveals loci linked to human variation in LINE-1- and Alu-insertion numbers"

### Figure S1

Supplementary Figure 1

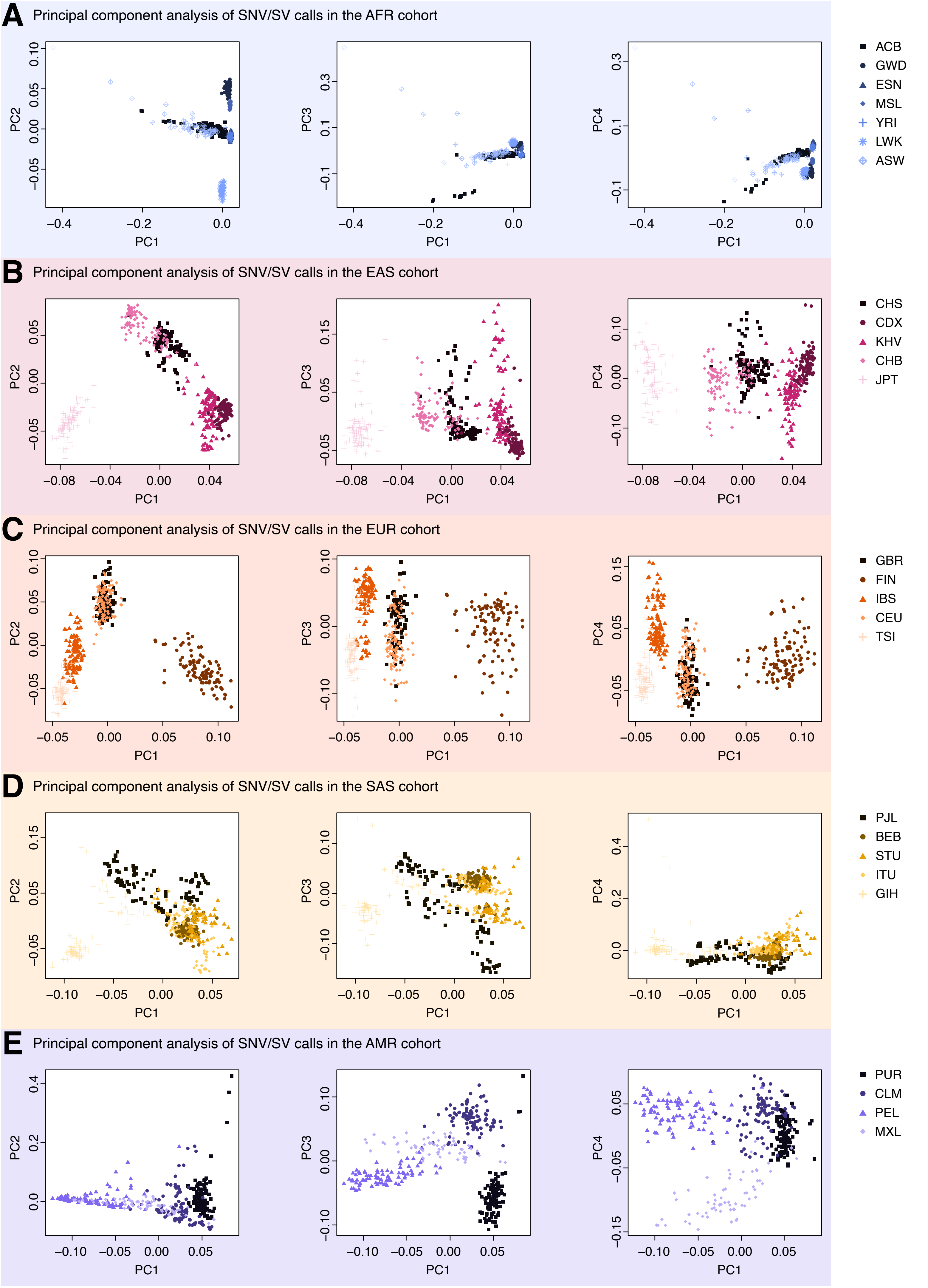

### Figure S2

Supplementary Figure 2

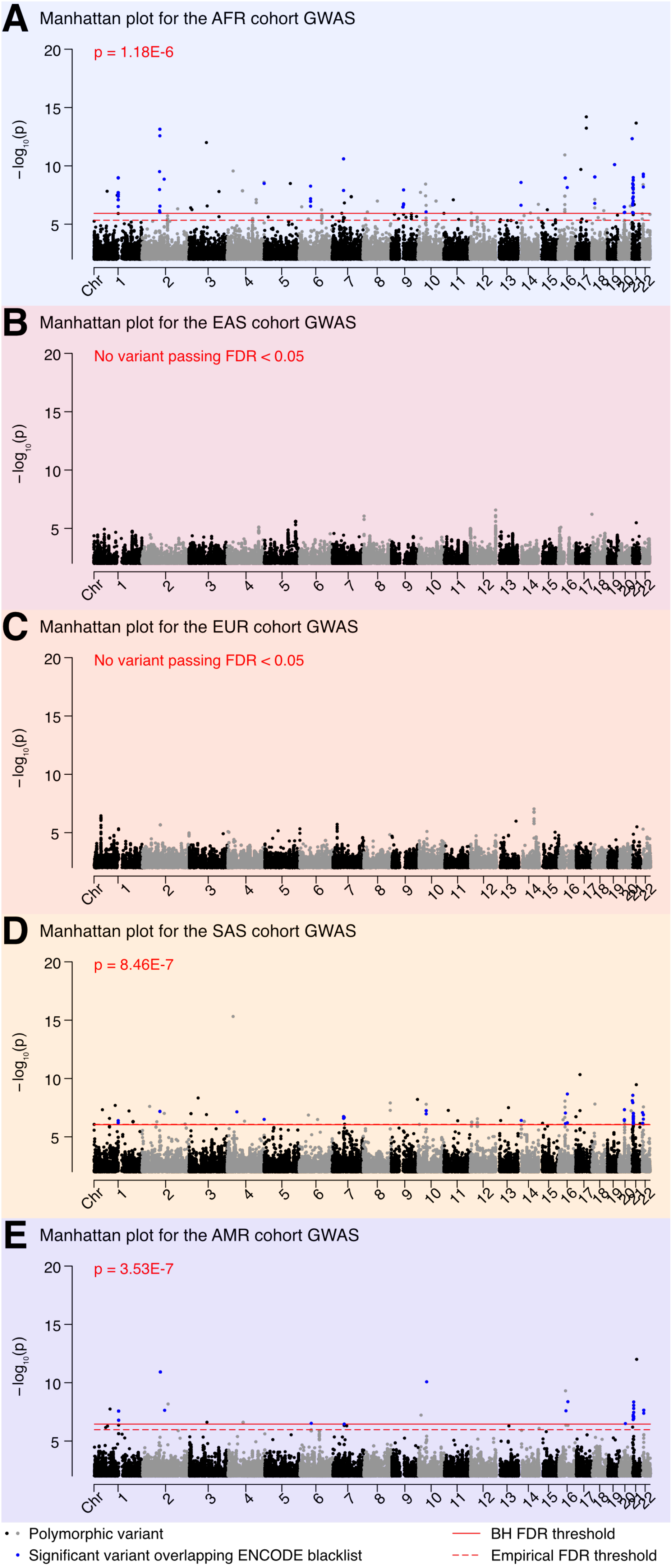

### Figure S3

#### Supplementary Figure 3

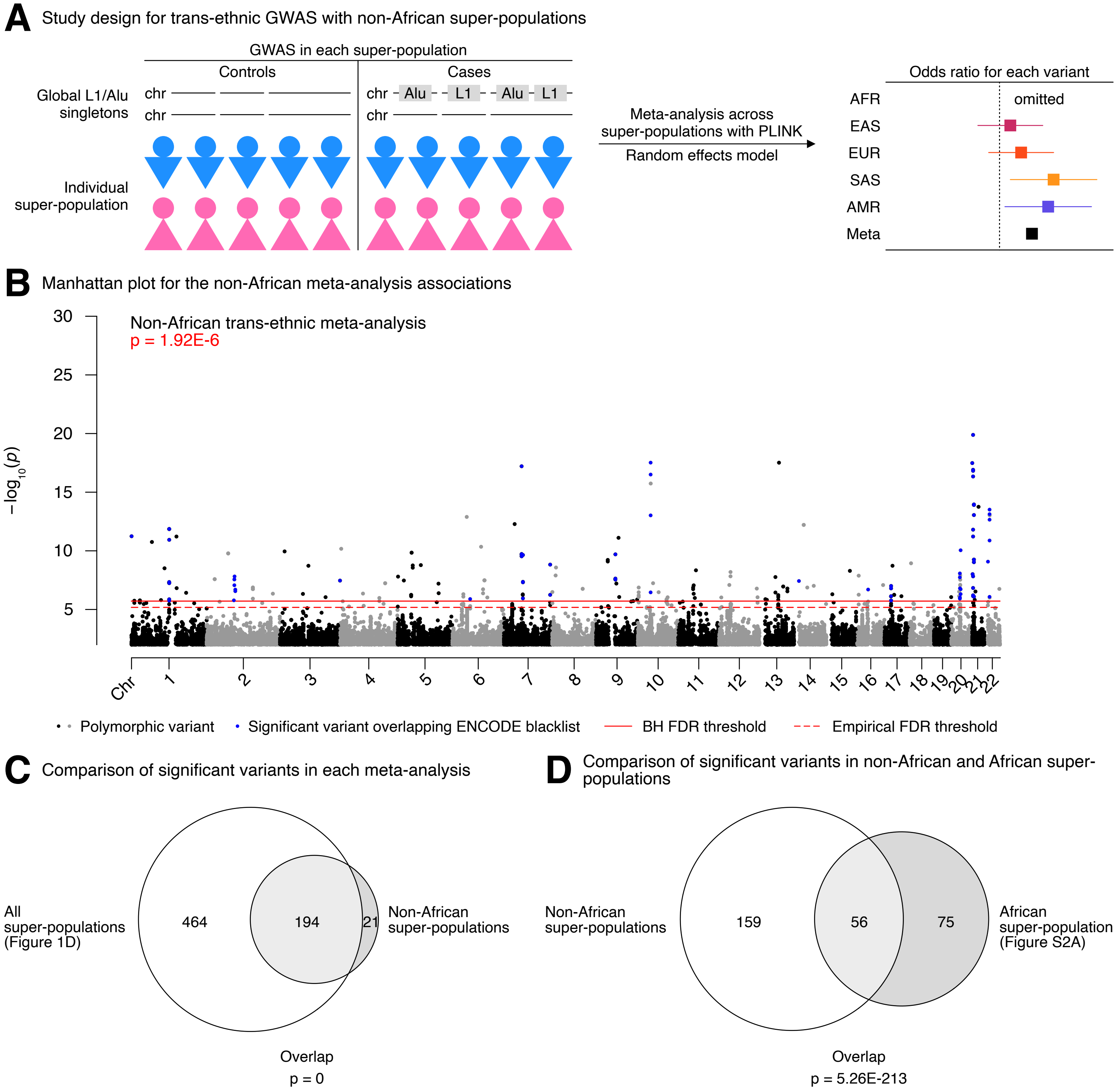

### Figure S4

Supplementary Figure 4

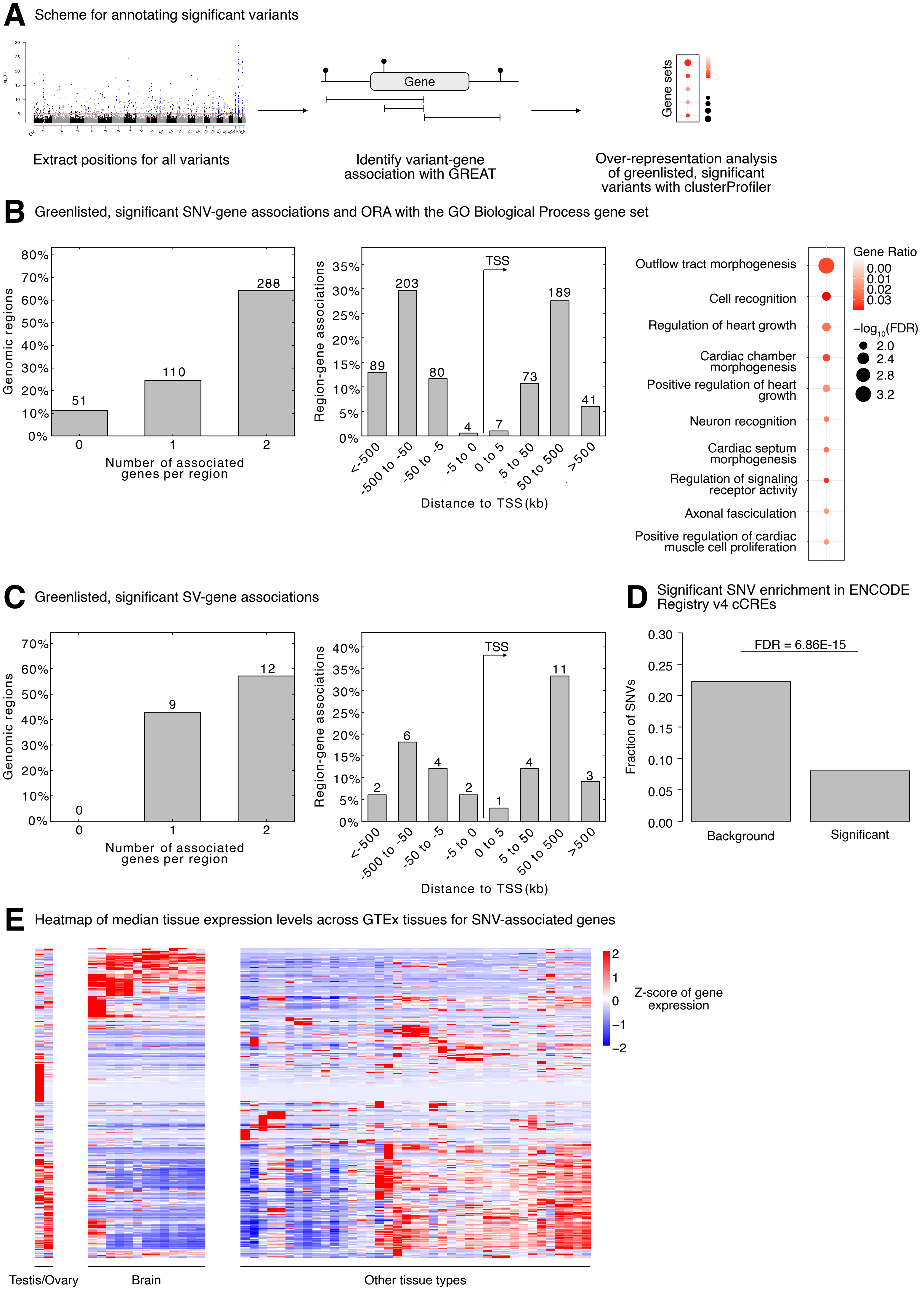

### Figure S5

Supplementary Figure 5

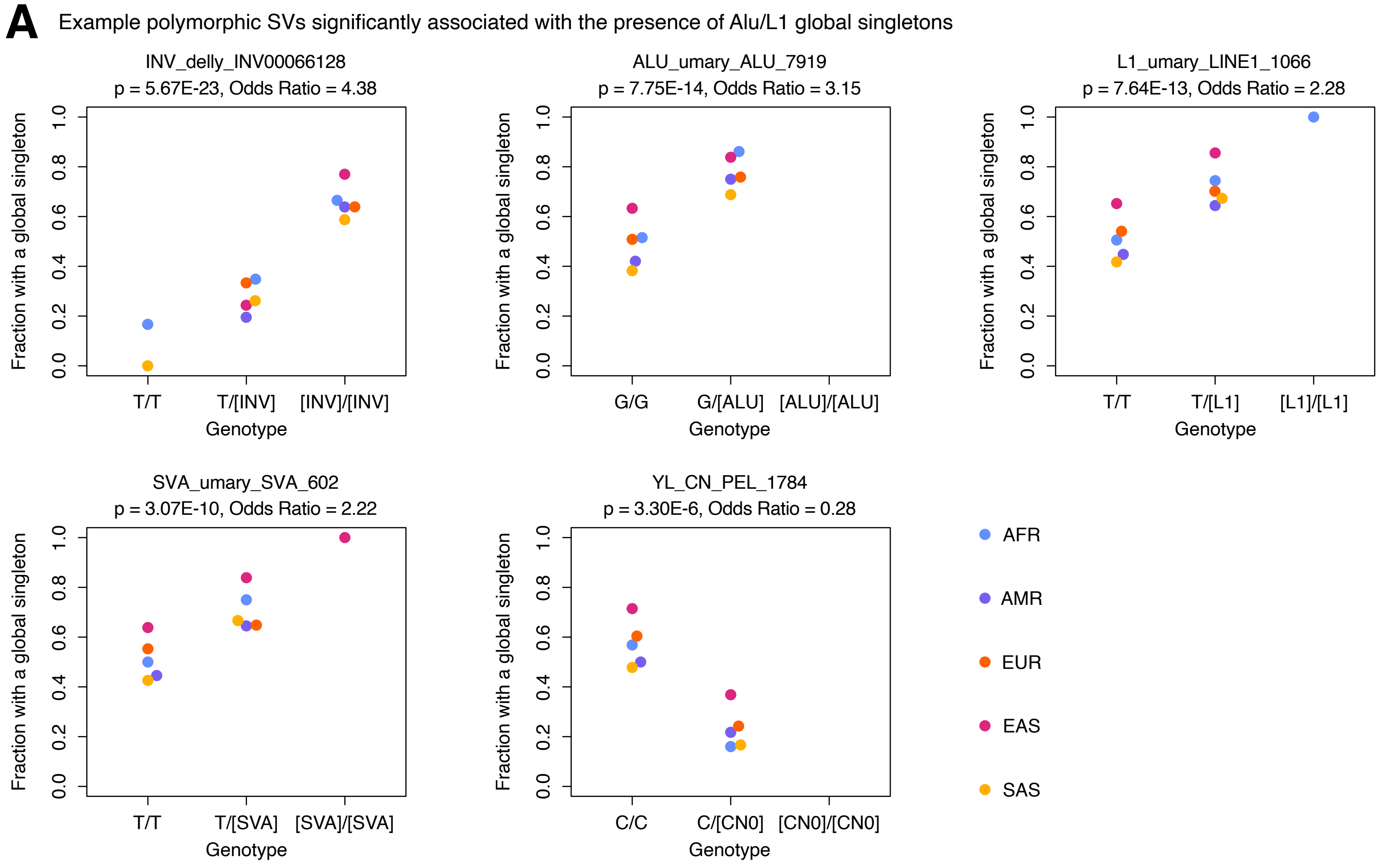
